# Strong evolutionary convergence of receptor-binding protein spike between COVID-19 and SARS-related coronaviruses

**DOI:** 10.1101/2020.03.04.975995

**Authors:** Yonghua Wu

**Affiliations:** School of Life Sciences, Northeast Normal University, 5268 Renmin Street, Changchun, 130024, China

## Abstract

Coronavirus Disease 2019 (COVID-19) and severe acute respiratory syndrome (SARS)-related coronaviruses (e.g., 2019-nCoV and SARS-CoV) are phylogenetically distantly related, but both are capable of infecting human hosts via the same receptor, angiotensin-converting enzyme 2, and cause similar clinical and pathological features, suggesting their phenotypic convergence. Yet, the molecular basis that underlies their phenotypic convergence remains unknown. Here, we used a recently developed molecular phyloecological approach to examine the molecular basis leading to their phenotypic convergence. Our genome-level analyses show that the spike protein, which is responsible for receptor binding, has undergone significant Darwinian selection along the branches related to 2019-nCoV and SARS-CoV. Further examination shows an unusually high proportion of evolutionary convergent amino acid sites in the receptor binding domain (RBD) of the spike protein between COVID-19 and SARS-related CoV clades, leading to the phylogenetic uniting of their RBD protein sequences. In addition to the spike protein, we also find the evolutionary convergence of its partner protein, *ORF3a*, suggesting their possible co-evolutionary convergence. Our results demonstrate a strong adaptive evolutionary convergence between COVID-19 and SARS-related CoV, possibly facilitating their adaptation to similar or identical receptors. Finally, it should be noted that many observed bat SARS-like CoVs that have an evolutionary convergent RBD sequence with 2019-nCoV and SARS-CoV may be pre-adapted to human host receptor ACE2, and hence would be potential new coronavirus sources to infect humans in the future.

## Introduction

The 2019 novel coronavirus (2019-nCoV, also called severe acute respiratory syndrome (SARS)-CoV-2) has caused the current outbreak of coronavirus disease (COVID-19), which has emerged as a serious public health concern. The clinical and pathological features caused by 2019-nCoV resemble those seen in SARS^1–3^, which is caused by SARS coronavirus (SARS-CoV). Both 2019-nCoV and SARS-CoV have been determined to be of bat origin, with possible intermediate hosts prior to infecting humans^4,5^. Phylogenetic studies have shown that 2019-nCoV and SARS-CoV belong to the subgenus *Sarbecovirus*, but they are distantly related^5–8^, with a sequence identity of 79.6% at the whole-genome level^5^. One recent study showed that 2019-nCoV is more similar to a bat coronavirus (RaTG13), with a sequence identity of 96.2% at the whole-genome level, than many other coronaviruses from different hosts, suggesting a phylogenetic affinity of 2019-nCoV to bat coronavirus compared with SARS-CoV^5^. Despite their relatively distant phylogenetic relationships, 2019-nCoV and SARS-CoV are both known to be capable of infecting humans using the same cell receptor, angiotensin-converting enzyme 2 (ACE2)^5,6,9,10^, and their protein structures of receptor-binding protein spike (S) are found to be highly similar to each other^10,11^, suggesting their phenotypic convergence. The spike protein is responsible for receptor binding and membrane fusion, and it is important for host tropism and transmission capacity^6^. The spike protein of coronaviruses comprises two subunits, S1 and S2. The S1 subunit contains a receptor-binding domain (RBD), which harbors a receptor-binding motif (RBM) to make complete contact with the receptor (i.e., ACE2)^12,13^.

Considering that 2019-nCoV and SARS-CoV are distantly related, but show high similarity in the RBD protein structure and can use the same cell receptor, ACE2^5–11^, an evolutionary convergence may have occurred between them. In the present study, we employ a recently developed molecular phyloecological approach^14–16^, which uses a comparative phylogenetic analysis of functional gene sequences to determine the genetic basis of phenotypic evolution, and we examine the possible molecular basis underlying the phenotypic convergence between 2019-nCoV and SARS-CoV. Our results reveal positive selection signals and evolutionary convergent amino acid sites of the spike protein in both 2019-nCoV and SARS-CoV and their related coronaviruses, providing new insights into understanding the evolutionary origin of their phenotypic convergence.

## Results and discussion

We used likelihood ratio tests based on the branch and branch-site models implemented in the codeml program of PAML^17^ to examine the possible Darwinian selection of all 11 genes annotated in the 2019-nCoV genome (NC_045512). Positively selected genes (PSGs) were found by the branch-site model (Table 1), independent of the initial value variation of parameters (kappa and ω). Specifically, for the branches leading to 2019-nCoV and SARS-CoV, no PSGs were found, nor were PSGs found along the branches leading to their sister coronaviruses. However, we detected PSGs along the ancestral branch (branch C) of 2019-nCoV and its sister taxon, RaTG13, as well as along the ancestral branch (branch K) of SARS-CoV and its sister taxa, WIV16 and Rs4231 (Table 1, Fig. 1). For branch C, three PSGs (*S*, *Orf1ab* and *N*) were found, and for branch K, only one gene (*S*) was found to be under positive selection (Table 1). *Orf1ab* encodes replicase and *N* encodes nucleocapsid^18^. Intriguingly, the *S* gene was subject to Darwinian selection in both branches, C and K. This gene encodes the spike protein, which mediates receptor binding and membrane fusion^6^. The finding of Darwinian selection on the spike protein may suggest its adaptive evolution to the host receptors. To further examine the possible adaptive evolution of 2019-nCoV and SARS-CoV to a human host, we used RELAX^19^ to analyze the relative selection intensity change of the 11 genes along the branches leading to 2019-nCoV and SARS-CoV compared with their most recent ancestors, branches C and K, respectively (Tables S1-2). Among the 11 genes examined, gene *S* along the 2019-nCoV branch exhibited a significant selection intensification signal (K = 30.54, *p* = 0.000, Table S1, Fig. S1), and this remained robust in four independent runs. We also found that the *ORF6* gene showed slight selection intensification (K = 1.81, *p* = 0.000) along the SARS-CoV branch (Table S2), while its statistical significance only received two supports among five independent runs. For the selectively intensified gene *S* along the 2019-nCoV branch, 0.46% of the amino acid sites (about five amino acids) were under positive selection, while most sites (86.63%) were under purification selection (Table S1). This may suggest that 2019-nCoV was subject to an adaptive evolution during its adaption to possible intermediate and/or human hosts.

**Fig. 1.**
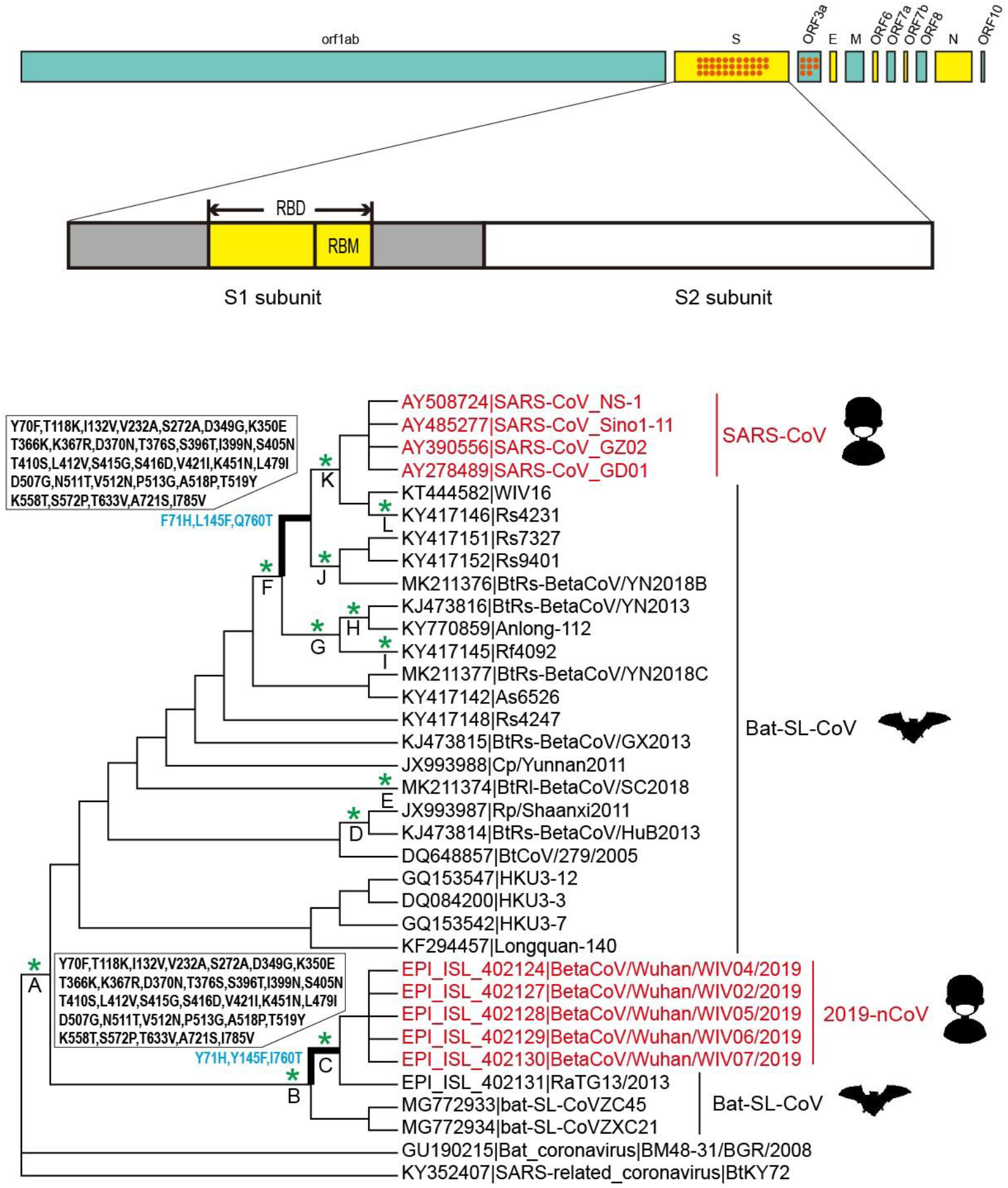
The phylogeny of subgenus *Sarbecovirus* and 11 genes used in this study. The coronavirus phylogeny follows two published studies^5,6^. 2019-nCoV, 2019 novel coronavirus; SARS-CoV, severe acute respiratory syndrome coronavirus; Bat-SL-CoV, bat-derived severe acute respiratory syndrome (SARS)-like coronaviruses. Genomic organization and the 11 genes annotated in the reference genome of 2019-nCoV (NC_045512) are shown. Red dots represent the numbers of identical or nearly identical amino acid sites found in genes, *S* and *ORF3a*, which are shared between SARS-related CoV and COVID-19-related CoV, but are completely or nearly completely distinct from those of their phylogenetic intermediates. The spike (S) protein structure follows one previous study^12^, and its receptor-binding domain (RBD) and receptor-binding motif (RBM) are highlighted. * above branches and their corresponding capital letters (A-L) denote the branches with positive selection signals found in the *S* gene. Two branches (bold) indicate two evolutionary convergent branches with three convergent amino acid substitutions (blue) and 32 shared parallel amino acid substitutions of spike protein.

**Table 1.** Positively selected genes identified based on the branch-site model. Only the ω values of the foreground branches are shown. Positive selection sites located in subunit 1 of gene *S* are shown in grey. Underlining shows positive selected sites located within RBD. Dashes (-) shows alignment gaps in the reference sequence.

| Branch/Gene | Parameter estimates | 2ΔL | df | p-value | Positive selection sites |
| --- | --- | --- | --- | --- | --- |
| **Branch C** |  |  |  |  |  |
| *S* | $p_0 = 0.899$ $p_1 = 0.067$ $p_{2a} = 0.030$ $p_{2b} = 0.002$<br>$\omega_0 = 0.029$ $\omega_1 = 1.000$ $\omega_{2a} = 27.064$ $\omega_{2b} = 27.064$ | 18.76 | 1 | 1.475E-05 | <u>19Y,67N,90N,187K,210M,289E,294S,351V</u><br><u>393G,399Y,420K,447T,463F,513T,539Q</u><br><u>551T,570-,577Q,615L,760A,770A</u> |
| *Orf1ab* | $p_0 = 0.944$ $p_1 = 0.050$ $p_{2a} = 0.004$ $p_{2b} = 0.000$<br>$\omega_0 = 0.029$ $\omega_1 = 1.000$ $\omega_{2a} = 206.003$ $\omega_{2b} = 206.003$ | 56.45 | 1 | 5.756E-14 | 154I,1756G,3909G,3918N,4019H,4222M,4228S<br>4287L,4298L,4319V,4446V,4478I,4486L,4502C<br>4864S,4960H,5882S,5930N,6128H,6191S,6323I<br>6396A,6428S,6436K,6488S |
| *N* | $p_0 = 0.914$ $p_1 = 0.080$ $p_{2a} = 0.005$ $p_{2b} = 0.000$<br>$\omega_0 = 0.037$ $\omega_1 = 1.000$ $\omega_{2a} = 122.692$ $\omega_{2b} = 122.692$ | 5.15 | 1 | 0.023 | 26D,104E,129E,219T,236V,336H,347N,415G |
| **Branch K** |  |  |  |  |  |
| *S* | $p_0 = 0.924$ $p_1 = 0.069$ $p_{2a} = 0.004$ $p_{2b} = 0.000$<br>$\omega_0 = 0.030$ $\omega_1 = 1.000$ $\omega_{2a} = 999.000$ $\omega_{2b} = 999.000$ | 7.29 | 1 | 0.006 | <u>11D,19Y,150N,390P</u> |

2 $\Delta$ L: twice difference of likelihood values between two nested models; df: degrees of freedom; proportion of sites and their corresponding $\omega$ values in four site classes ( $p_0$ , $p_1$ , $p_{2a}$ and $p_{2b}$ ) are shown.

Given the positive selection of the *S* gene in both branches, C and K, we further examined its positive selection signals using branch-site model along all other main branches (Fig. 1) to test whether the positive selection uniquely occurred along the branches related to 2019-nCoV and SARS-CoV. Our results showed positive selection signals along 10 out of 45 branches examined (Fig. 1, Table S3). This may suggest that the *S* gene was widely subject to Darwinian selection in different coronavirus strains, indicating that it may be crucial for the successful survival of coronaviruses. Further analyses of positive selection sites showed that most positive selection sites among all 12 branches under positive selection were located within subunit 1 of *S* gene (Table 1, Table S3), which is used for receptor binding. This may suggest that there are strong selection pressures of different coronavirus strains for their own receptor binding.

Given the selection intensification of *S* gene in 2019-nCoV since its evolutionary divergence from RaTG13, we conducted comparative sequence analyses between the two. We found that there were more than 20 amino acid differences between them, and most of them were located within RBD, especially RBM (Fig. S2), suggesting a high variability of RBM. Given the importance of RBM for receptor binding, we further conducted a comparative sequence analysis among all the coronavirus strains studied to examine its variability. The results showed high sequence variability, with insertion and/or deletion and amino acid substitutions among the coronavirus strains studied (Fig. 2). Despite the high variability of RBM, strikingly, we found that SARS-CoV and its phylogenetic relatives—including WIV16 (KT444582), Rs4231 (KY417146), Rs7327 (KY417151), Rs9401 (KY417152), and BtRs-BetaCoV/YN2018B (MK211376), called SARS-related CoV here, shared many identical or nearly identical amino acids with their phylogenetically distant coronavirus strains, including 2019-nCoV and RaTG13, which we called COVID-19-related CoV (Fig. 2). These shared amino acids were clearly distinct from bat SARS-like CoV that were phylogenetic intermediates between them (Fig. 2). Further analyses showed that such identical amino acids shared between SARS-related CoV and COVID-19-related CoV were not restricted to RBM, but rather, they were scattered throughout the spike protein, with a total of 32 such sites, which were centered on RBD (28 sites in total, Fig. S3). To further examine whether such similarity occurred in other proteins, we analyzed all 11 genes studied among these coronaviruses, and we found that one additional gene, *ORF3a*, contained eight such sites (Fig. 1). The existence of these shared amino acids between SARS-related CoV and COVID-19-related CoV may suggest their high sequence similarity. In support of this, we reconstructed maximum likelihood and neighbor-joining phylogenies using full-length RBD protein sequences, and both showed that SARS-related CoV and COVID-19-related CoV were grouped in the same clade, with relatively high support, which is consistent with two previous studies^6,7^, at the same time, their phylogenetic intermediates were clustered in distinct clades (Fig. 3, Fig. S4). The phylogenetic uniting of SARS-related CoV and COVID-19-related CoV provide evidence of their high similarity of RBD protein sequences.

**Fig. 2.**
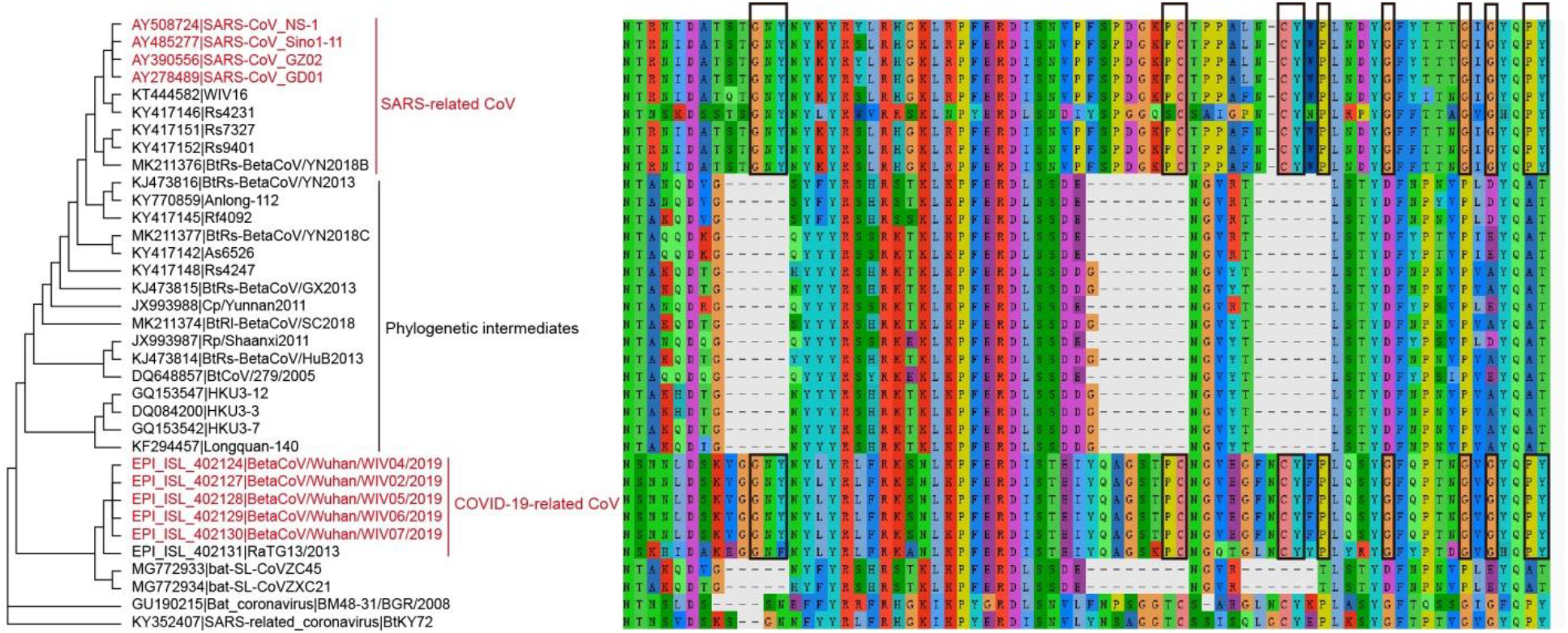
The identical or nearly identical RBM amino acid sites (rectangle) shared between SARS-related CoV and COVID-19-related CoV. These shared amino acids are distinct from that of their phylogenetic intermediates. The phylogeny is the same as in Fig. 1.

**Fig. 3.**
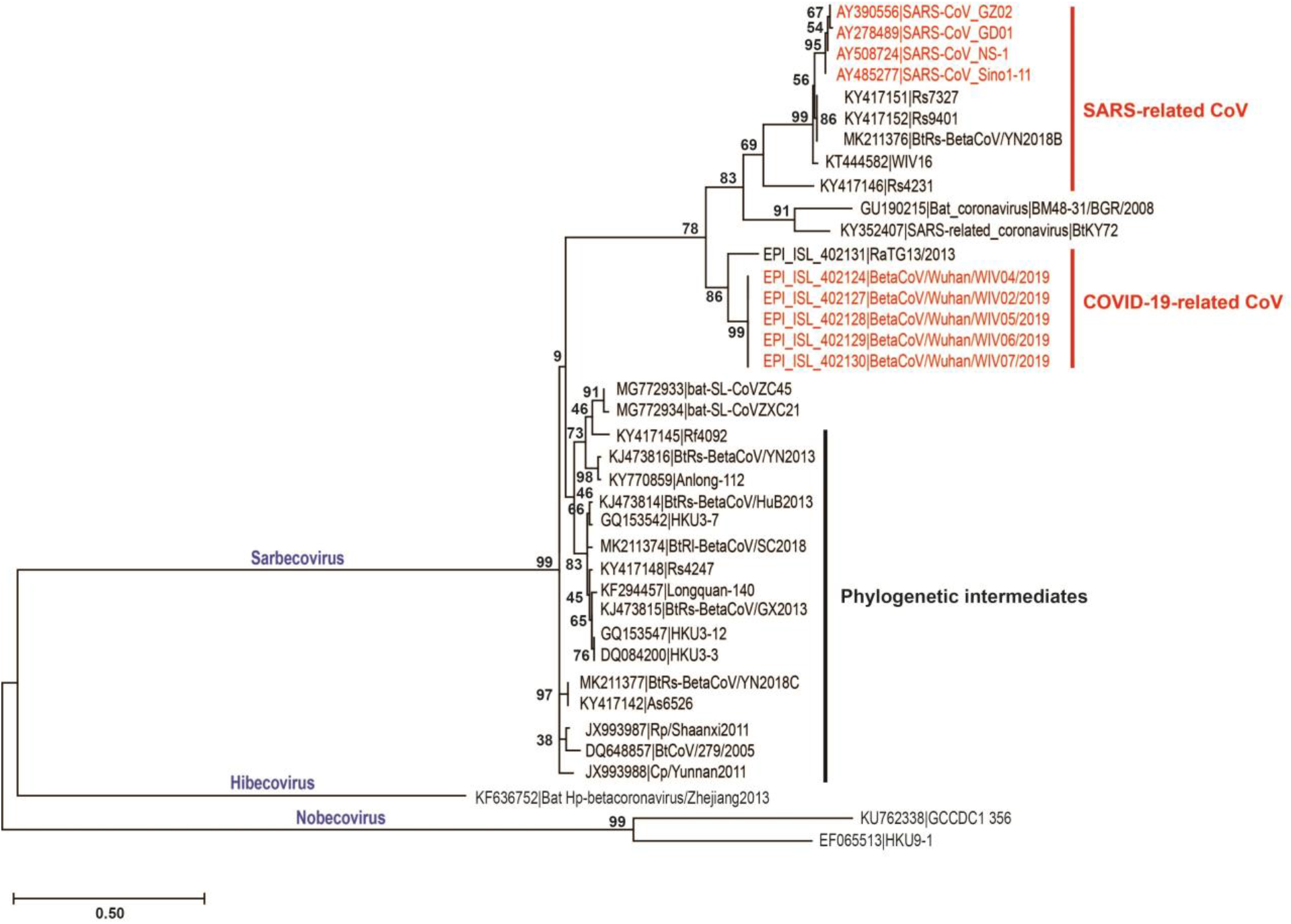
Maximum likelihood tree of the full-length amino acid sequence of RBD. The WAG+G amino acid substitution model is used. The tree has the highest log likelihood (−3406.88). The node supports are shown in number. *Hibecovirus* and *Nobecovirus* are used as outgroups.

Given their genome-level phylogenetic disparity, the high similarity of RBD protein sequences between SARS-related CoV and COVID-19-related CoV may suggest their evolutionary convergence in the spike protein. To test this possibility, we used an empirical Bayes approach in PAML^17^ to reconstruct ancestral amino acid sequences along internal nodes, and our results showed there were up to 35 evolutionary convergent sites, including 3 convergent and 32 parallel amino acid substitutions that were shared by two ancestral branches leading to SARS-related CoV and COVID-19-related CoV, respectively (Fig. 1). It should be noted that the 35 evolutionary convergent sites of spike protein were apparently underestimated, since those amino acid sites with alignment gaps were not considered by the approach used. Still, these 35 sites represented an unusually high incidence of evolutionary convergence sites, which have rarely been found in previous studies related to molecular convergent evolution^20–26^. This suggests a strong evolutionary convergence between SARS-related CoV and COVID-19-related CoV. On completion of our data analyses, one of the most recent studies showed that the RBD of the spike protein of Pangolin-CoV is nearly identical to that of 2019-nCoV, with only one amino acid difference^27^, suggesting that Pangolin-CoV also belongs to the clade of COVID-19-related CoV. It should be noted that the RBD sequences of two other coronavirus strains, BM48-31 (GU190215) and BtKY72 (KY352407), were grouped with SARS-related CoV and COVID-19-related CoV (Fig. 3, Fig. S4), suggesting an evolutionary convergence among them. In addition to these evolutionarily convergent coronavirus strains, intriguingly, we found the evidence of evolutionary convergence of the spike protein between the ancestral branch leading to SARS-related CoV and the ancestral branch of COVID-19-related CoV and its sister strains (CoVZC45 and CoVZXC21), which harbored 9 evolutionary convergent amino acid sites (Fig. S5). These results suggest that the spike protein may have been subjected to a successive evolutionary convergence among ancestral coronavirus strains leading to SARS-related CoV, COVID-19-related CoV and CoVZC45 and CoVZXC21.

Previous studies show that spike protein interacts tightly with a related protein *ORF3a* and they likely coevolved^28–30^. If their coevolution does occur, we may expect that the evolutionary convergence of the spike protein found may have led to the occurrence of the evolutionary convergence of *ORF3a* as well. To test this, we reconstructed ancestral amino acid sequences of *ORF3a* along internal nodes, and our results revealed 6 parallel amino acid substitutions shared between the ancestral branch leading to SARS-related CoV and the ancestral branch of COVID-19-related CoV and its sister strains (CoVZC45 and CoVZXC21) (Fig. S5). And we also detected a parallel amino acid substitution of *ORF3a* between the two ancestral branches leading to SARS-related CoV and COVID-19-related CoV (Fig. S6). Considering that these evolutionary convergent branches of *ORF3a* also showed evolutionary convergence in spike protein as mentioned above (Fig. 1, Fig. S5), it may suggest that spike protein and its partner protein *ORF3a* may have been subjected to a co-evolutionary convergence.

Evolutionary convergence may occur by chance or by Darwinian selection. Our results showed that the evolutionary convergent sites found in this study were mainly restricted to two genes (*S* and *ORF3a*, Fig. 1), and in particular, they were centered within the RBD of the *S* gene. This biased distribution of evolutionary convergent sites is difficult to explain according to chance; rather, Darwinian selection would be favored as a plausible explanation. In support of this, we used CONVERG2^31^ to evaluate the probability of the occurrence of our observed convergent sites of spike protein between the two ancestral branches leading to SARS-related CoV and COVID-19-related CoV, and the results showed high statistical significance (*p* = 0.000000), regardless of whether the JTT model or Poisson model was used. This result apparently rejects chance or neutral evolution as a possible explanation; rather, it indicates a predominately strong Darwinian selection. Moreover, we observed an apparently accelerated evolution of RBD of SARS-related CoV and COVID-19-related CoV related to their phylogenetic intermediates (Fig. 3), and we detected a significant Darwinian selection of the *S* gene along two branches (branches C and K) of SARS-related CoV and COVID-19-related CoV (Fig. 1, Table 1). These lines of evidence may strongly support the evolutionary convergence found in this study as a result of adaptive evolution. Regarding the possible adaptive evolutionary convergence, previously proposed causes, such as gene duplication and horizontal gene transfer^21,32,33^, are less likely because only single-copy *S* genes were found in all 35 genomes examined and evolutionary convergent sites presented an apparently biased distribution pattern. Parallel and/convergent evolution, which occur through point mutation, could contribute to our observed evolutionary convergence, but it could not account for the unusually high incidence of convergent sites observed in this study, representing a rare finding in previous studies^20–26^. Recent studies have shown a relatively high likelihood of occurrence of homologous recombination in spike protein^7,34,35^, and especially, it is considered that the RBD of 2019-nCov may be derived from a recombination event between that of human SARS-CoV and another (unsampled) SARS-like CoV^35^. If this is the case, the homologous recombination, if any, may have occurred between the ancestors (branches C and K) of SARS-related CoV and COVID-19-related CoV, accounting for their unusually high incidence of convergent sites observed in this study.

Given the strong evolutionary convergence of RBD of spike protein between the two clades, COVID-19-related CoV and SARS-related CoV, the coronaviruses of the two clades may have more likely adapted to similar or the same receptor. To date, 2019-nCoV and SARS-CoV have been known to be capable of using the ACE2 receptor in human host^5,6,9,10^, but the receptors of their phylogenetic relatives from the two clades, COVID-19-related CoV and SARS-related CoV, are less clear^4,5,36,37^. Regarding the bat SARS-like CoV, Rs4231 and Rs7327 are known to be able to use human ACE2 receptor^37^, while WIV16 is capable of using the ACE2 receptor from humans, civets and Chinese horseshoe bats (*Rhinolophus sinicus*)^4^. The receptors of Rs9401, BtRs-BetaCoV/YN2018B, and BatCoV RaTG13 remain to be explored. Further studies on the receptors of these bat SARS-like CoVs in their natural reservoirs are badly needed to determine whether ACE2 or other candidates, if any, represent their shared cell receptor, leading to their strong evolutionary convergence of spike protein.

Our molecular phyloecological study demonstrates that spike protein shows significant Darwinian selection along two ancestral branches related to SARS-CoV and 2019-nCoV, suggesting their adaptive evolution to recognizing their own cell receptors. Comparative sequence and phylogenetic analyses indicate a high similarity of RBD sequences of spike protein between SARS-related CoV and COVID-19-related CoV. Subsequent ancestral sequence reconstruction and convergent evolution analyses reveal an unusually high incidence of parallel and convergent amino acid substitutions between them, suggesting an extremely strong adaptive evolutionary convergence in spike protein. In addition to spike protein, we also found evolutionary convergence of its partner protein, *ORF3a*, suggesting their possible co-evolutionary convergence. Finally, considering that SARS-CoV and 2019-nCoV have posed serious concerns to public health and safety, it should be noted that many other bat SARS-like CoV strains that were evolutionarily convergent with SARS-CoV and 2019-nCoV recognized in this study may be potential novel coronaviruses to infect humans in the future.

## Materials and method

### Taxa and sequences

We used 35 coronavirus strain genomes of the subgenus *Sarbecovirus* based on two published studies^5,6^, including 2019-nCoV, SARS-CoV, and their phylogenetic relative, bat SARS-like CoV (please see Fig. 1 for details). For all these coronavirus strains, we downloaded their full-length genome sequences from GenBank except for five 2019-nCoV and RaTG13 strains, which were downloaded from GISAID. We aligned these genome sequences using MAFFT (https://mafft.cbrc.jp/alignment/server/). The coding sequences of 11 genes (Fig. 1) annotated in the genome of 2019-nCoV (NC_045512) were used as a reference sequence to obtain the homologous gene sequences of the 35 coronavirus strains. We aligned these homologous gene sequences using the online software webPRANK (http://www.ebi.ac.uk/goldman-srv/webprank/)^38^, which is considered to create a more reliable alignment to decrease false-positive results in positive selection analyses^39^.

### Adaptive evolution analyses

We employed the branch and branch-site models implemented in the codeml program of PAML^17^ to examine the adaptive evolution of our focal genes. For this, a codon-based maximum-likelihood method was used to estimate the ratio of non-synonymous to synonymous substitutions per site (dN/dS or ω), and likelihood ratio tests (LRTs) were used to calculate statistical significance. A statistically significant value of ω > 1 suggests positive selection. Upon analysis, an unrooted taxon tree (Fig. 1) was constructed based on two published studies^5,6^. For branch model analysis, we used a two-rate branch model, and our focal branches were labelled as foreground branches, while others were treated as background branches. The two-rate branch model was compared with the one-rate branch model, which assumes a single ω value across the tree, to determine statistical significance. If a statistically significant value of ω > 1 in a foreground branch was detected, the two-ratio branch model was then compared with the two-ratio branch model with a constraint of ω = 1 to further determine whether the ω > 1 of the foreground branch was statistically significant. In addition to the branch model, we also used a branchsite model (Test 2) to detect positively selected sites for a particular branch. Test 2 compares a modified model A with its corresponding null model with a constraint of ω = 1 to determine the statistical significance. Positively selected sites were found using an empirical Bayes method. For result robustness, we evaluated the dependence of the signal of positive selection on parameter variation. For this, we used two different initial values of kappa (kappa = 0.5, 3.0) and of omega (ω = 0.5, 2.0), and eventually, several independent runs were conducted for each of the positively selected genes found.

### Selection intensity analyses

We analyzed relative selection intensity using the RELAX^19^ program, available from the Datamonkey webserver (http://test.datamonkey.org/relax). RELAX is a hypothesis testing framework, and it can be used to test whether selection strength has been relaxed or intensified along a certain branch or lineage. For analyses, RELAX calculates a selection intensity parameter value (k), and k>1 shows an intensified selection, while k<1 indicates a relaxed selection, assuming a priori partitioning of the test branches and reference branches. We would expect that an intensified selection shows ω categories away from neutrality (ω = 1), while a relaxed selection is expected to show ω categories converging to neutrality (ω = 1). Statistical significance was evaluated by LRT by comparing an alternative model with a null model. The null model assumes k = 1 and the same ω distribution for both test and reference branches, while the alternative model assumes that k is a free parameter, and the test and reference branches may have different ω distributions.

### Phylogenetic analyses

We reconstructed an maximum likelihood (ML) tree and neighbor-joining (NJ) tree using MEGA X^40^. For ML analyses, the WAG+ G model was selected as the best amino acid substitution model according to the Bayesian information criterion. All amino acid sites with an alignment gap were included for analyses. For NJ analyses, JTT+G was selected as the best model. For the ML and NJ analyses, the bootstrap value was set to 1, 000. Other parameters were used as defaults in the program.

### Ancestral sequence reconstruction

We used the amino acid-based marginal reconstruction implemented in the empirical Bayes approach in PAML^17^ for ancestral sequence reconstruction. In the analyses, the character was assigned to a single interior node and the character with the highest posterior probability was used as the best reconstruction. We used two different amino acid substitution models, JTT and Poisson, to examine the consistency of our results. The JTT model assumes different substitution rates of different amino acids, while the Poisson model assumes the same substitution rate of all amino acids. For the analyses, we obtained the full-length spike protein sequences of our focal coronavirus strains and used their phylogeny, as given in Fig. 1. The amino acid substitutions along our focal branches were analyzed. The results based on the JTT and Poisson models were generally identical; for convenience, only the results based on the JTT model are shown.

### Convergent evolution analyses

We used the CONVERG2^31^ program to evaluate the probabilities that the observed convergent and parallel substitutions were due to random chance. A statistical significant *p*-value may suggest the observed evolutionary convergent sites are less likely attributable to random chance, but instead, favor Darwinian selection as a possible explanation. For the analyses, two different amino acid substitution models, JTT and Poisson, were used. The RBD amino acid sequences of our focal 35 coronavirus genomes were abstracted and aligned using CLUSTAL W^41^ program. The phylogenetic relationships among the coronavirus strains studied are given in Fig. 1.

## Acknowledgements

We thank Hui Wang for helping with the convergent evolution analyses. This research was supported by the National Natural Science Foundation of China (grant number, 31770401) and the Fundamental Research Funds for the Central Universities.

---

**Fig. S1.**
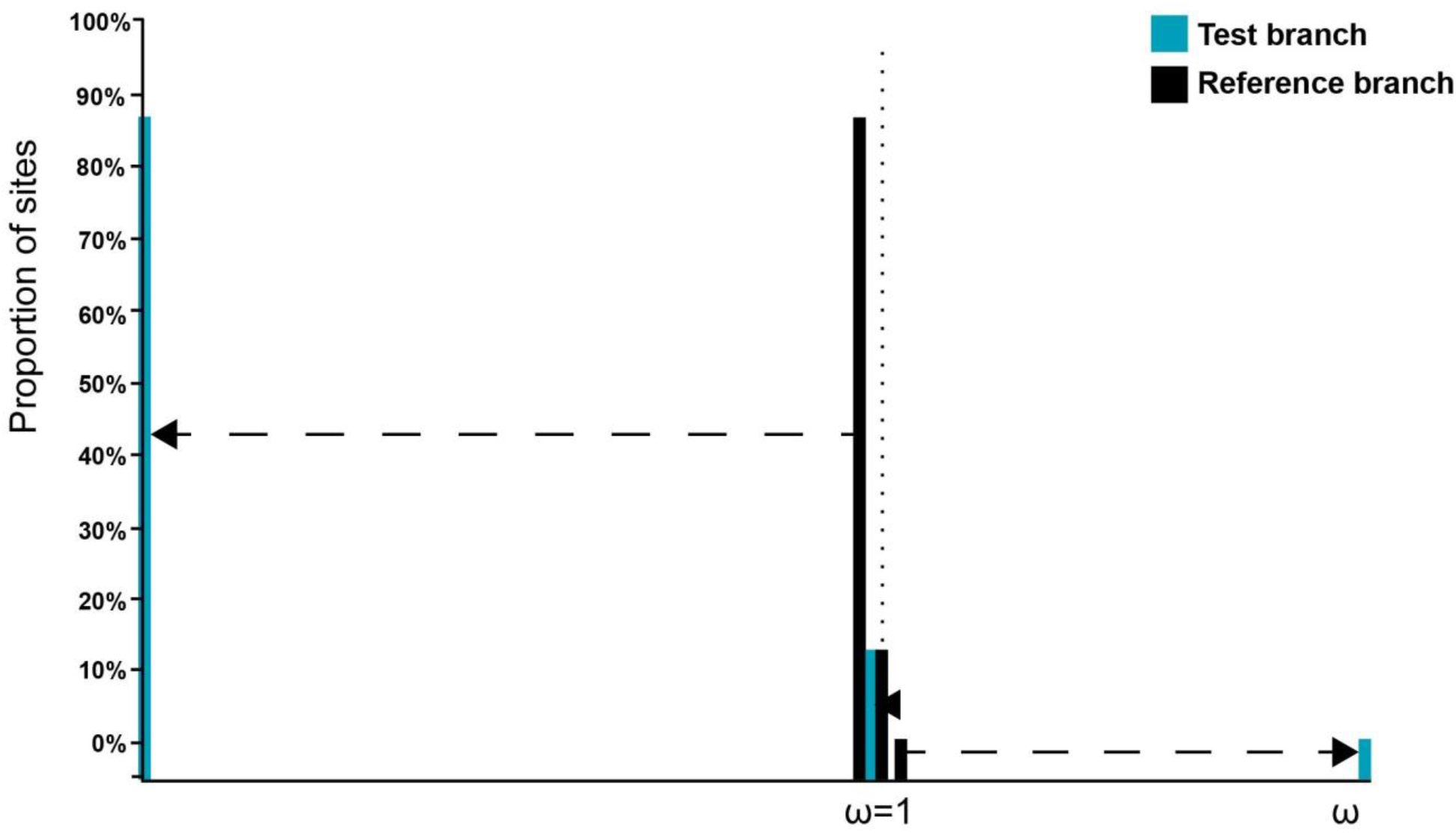
Selection intensity changes of gene *S* along the 2019-nCoV branch (test branch) compared with the common ancestral branch (reference branch) of 2019 n-CoV and RaTG13. The result shows that the ω categories of the test branch are apparently away from neutrality (ω = 1), indicating an intensified selection along the 2019-nCoV branch.

**Fig. S2.**
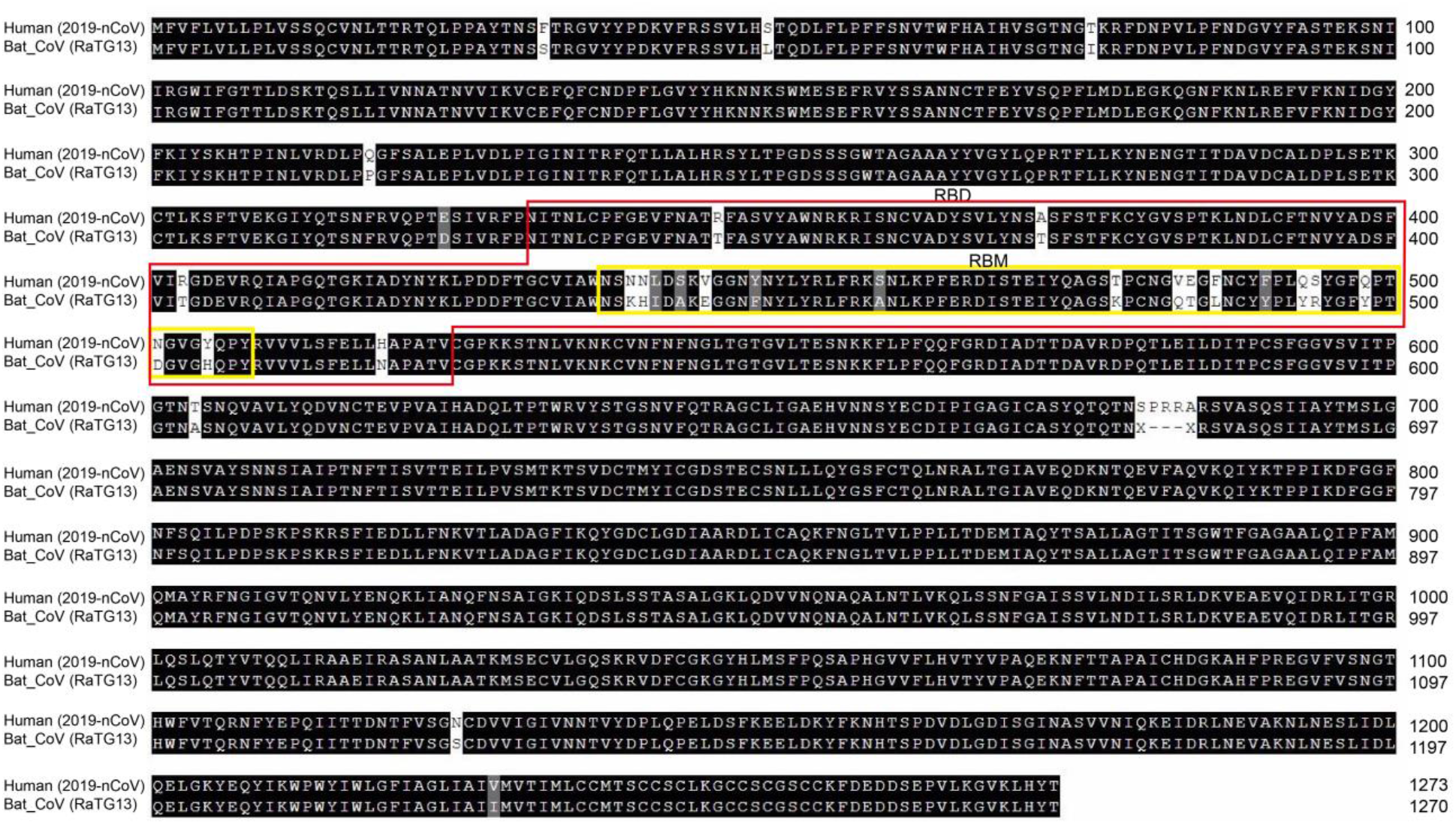
Amino acid variations of full-length spike protein sequences of 2019-nCoV and RaTG13. RBD, receptor-binding domain; RBM, receptor-binding motif.

**Fig. S3.**
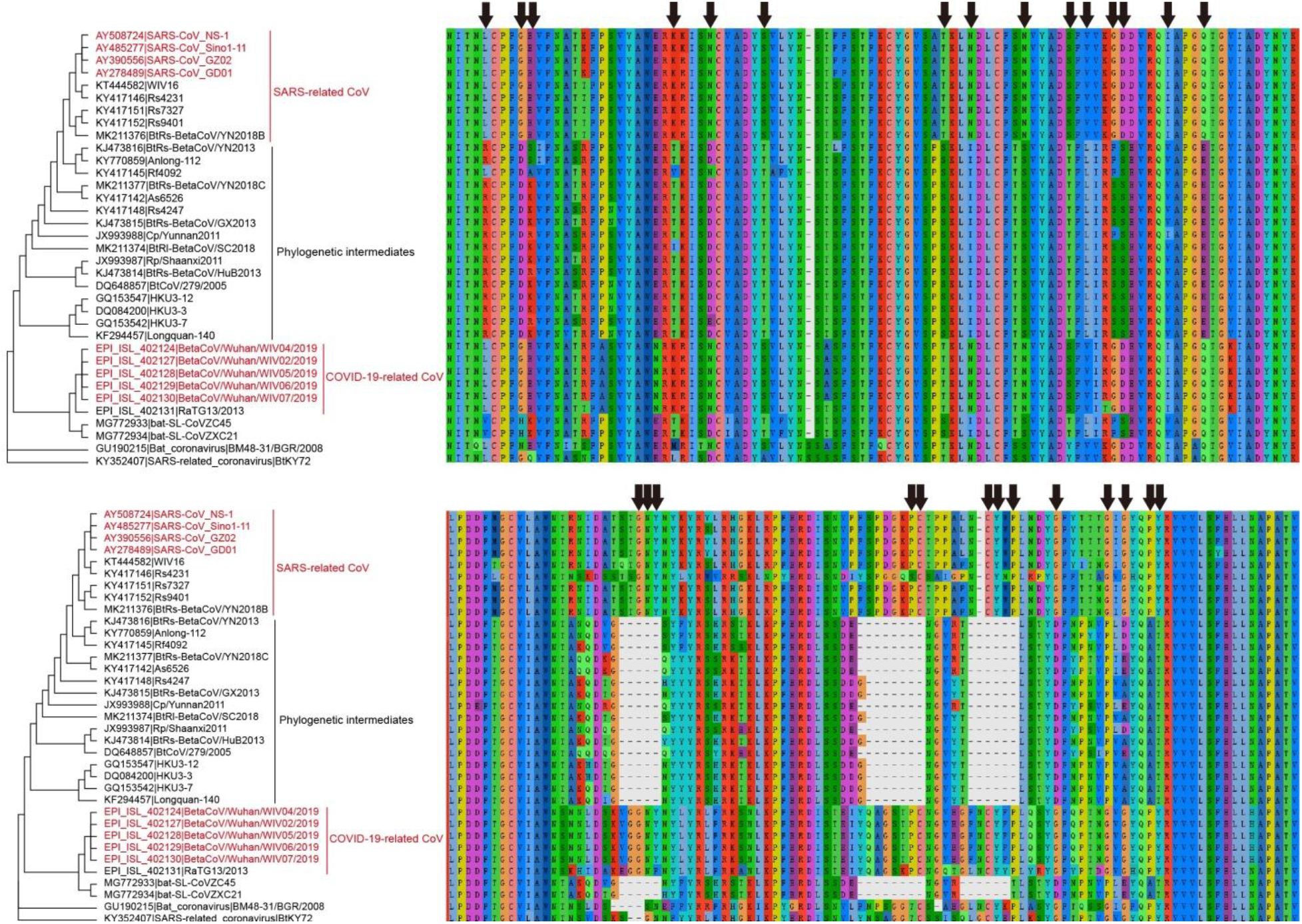
Twenty-eight identical or nearly identical RBD amino acid sites (arrows) shared between SARS-related CoV and COVID-19-related CoV. These shared amino acids are completely or nearly completely distinct from those of their phylogenetic intermediates. The phylogeny is the same as in Fig. 1.

**Fig. S4.**
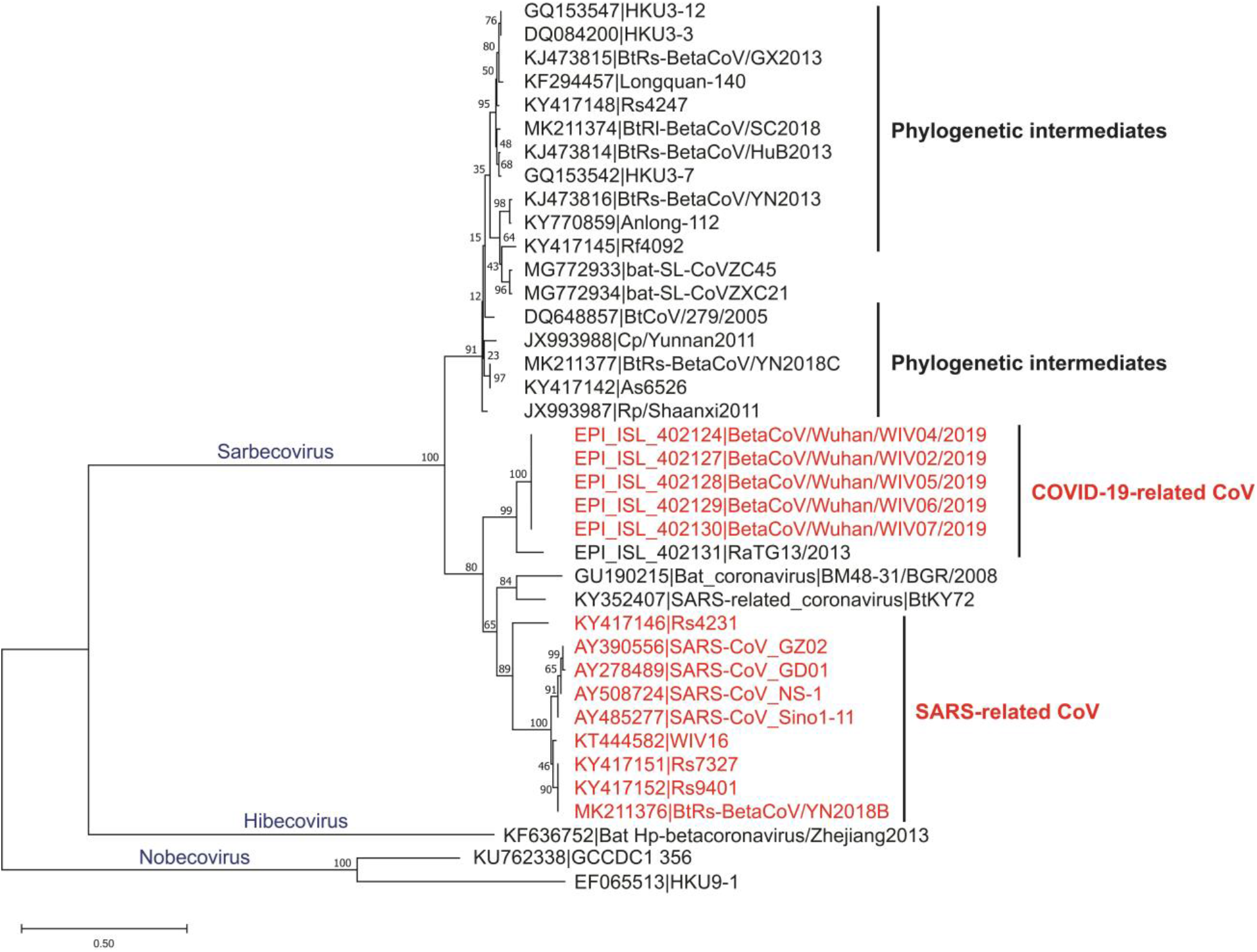
Neighbor-joining tree based on full-length amino acid sequence of RBD. The JTT+G model is used. The node supports are shown in numbers. *Hibecovirus* and *Nobecovirus* are used as outgroups.

**Fig. S5.**
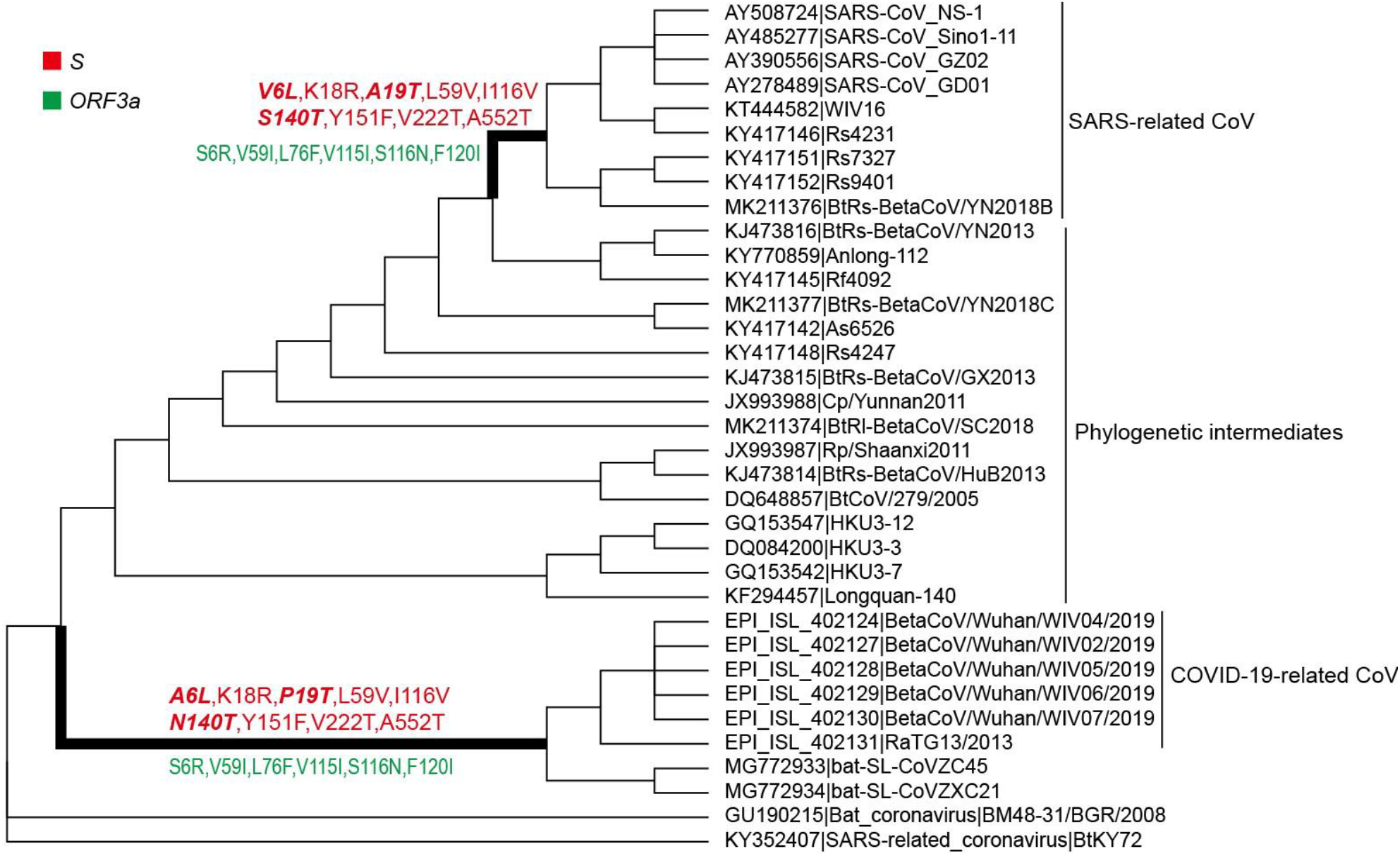
Convergent (bold italic) and parallel amino acid substitutions of genes *S* and *ORF3a*. The phylogeny is the same as in Fig. 1.

**Fig. S6.**
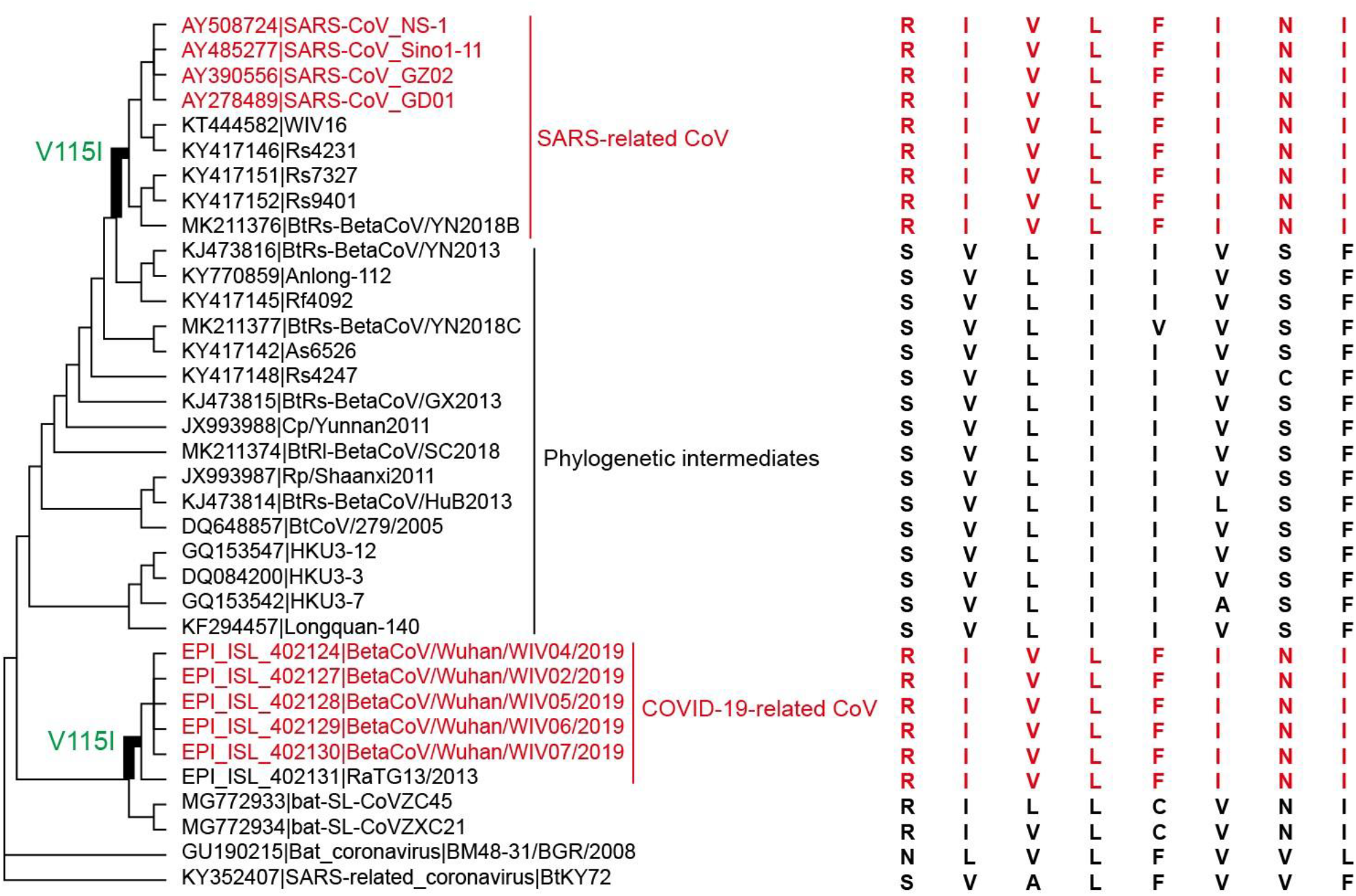
One parallel amino acid substitution of *ORF3a* protein shared by two ancestral branches (bold) leading to SARS-related CoV and COVID-19-related CoV. This parallel amino acid substitution is only supported as Poisson model was used. Eight identical amino acid sites of *ORF3a* protein shared between SARS-related CoV and COVID-19-related CoV are also shown. These shared amino acids are completely distinct from those of their phylogenetic intermediates. The phylogeny is the same as in Fig. 1.

**Table S1.**
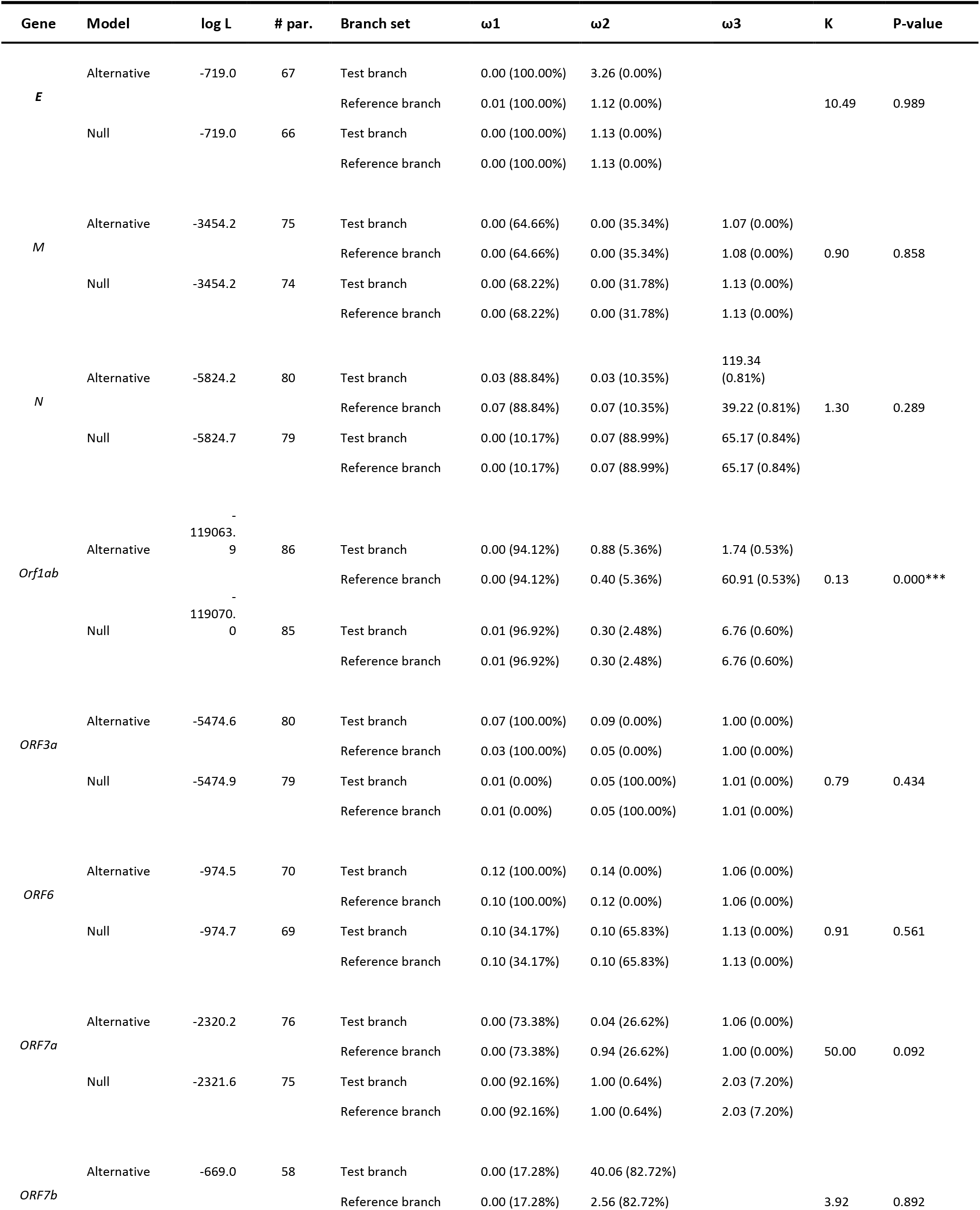
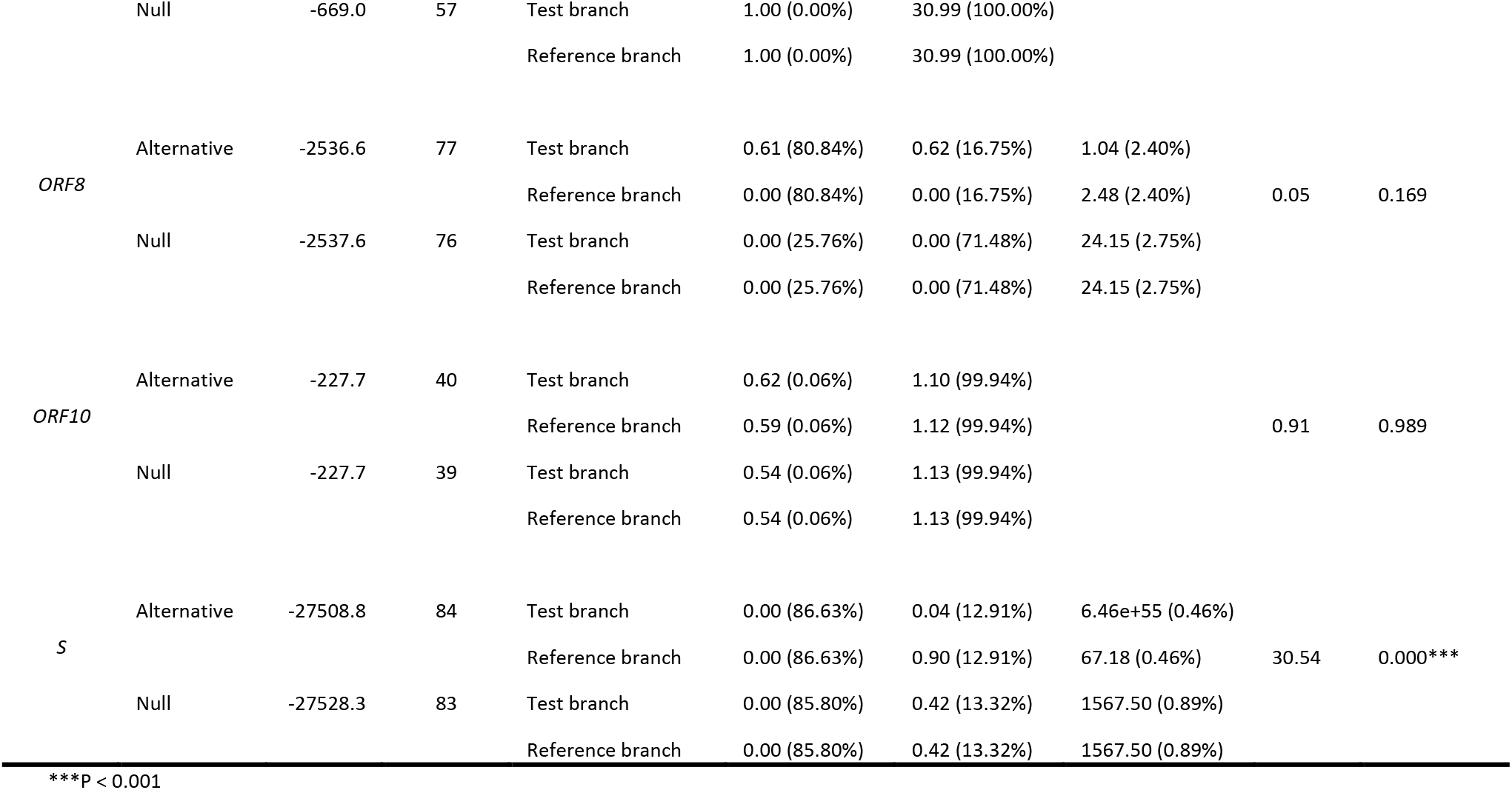
Selection intensity change of 11 genes along the 2019-nCoV branch (test branch) relative to the common ancestral branch (reference branch) of 2019-nCoV and RaTG13.

**Table S2.**
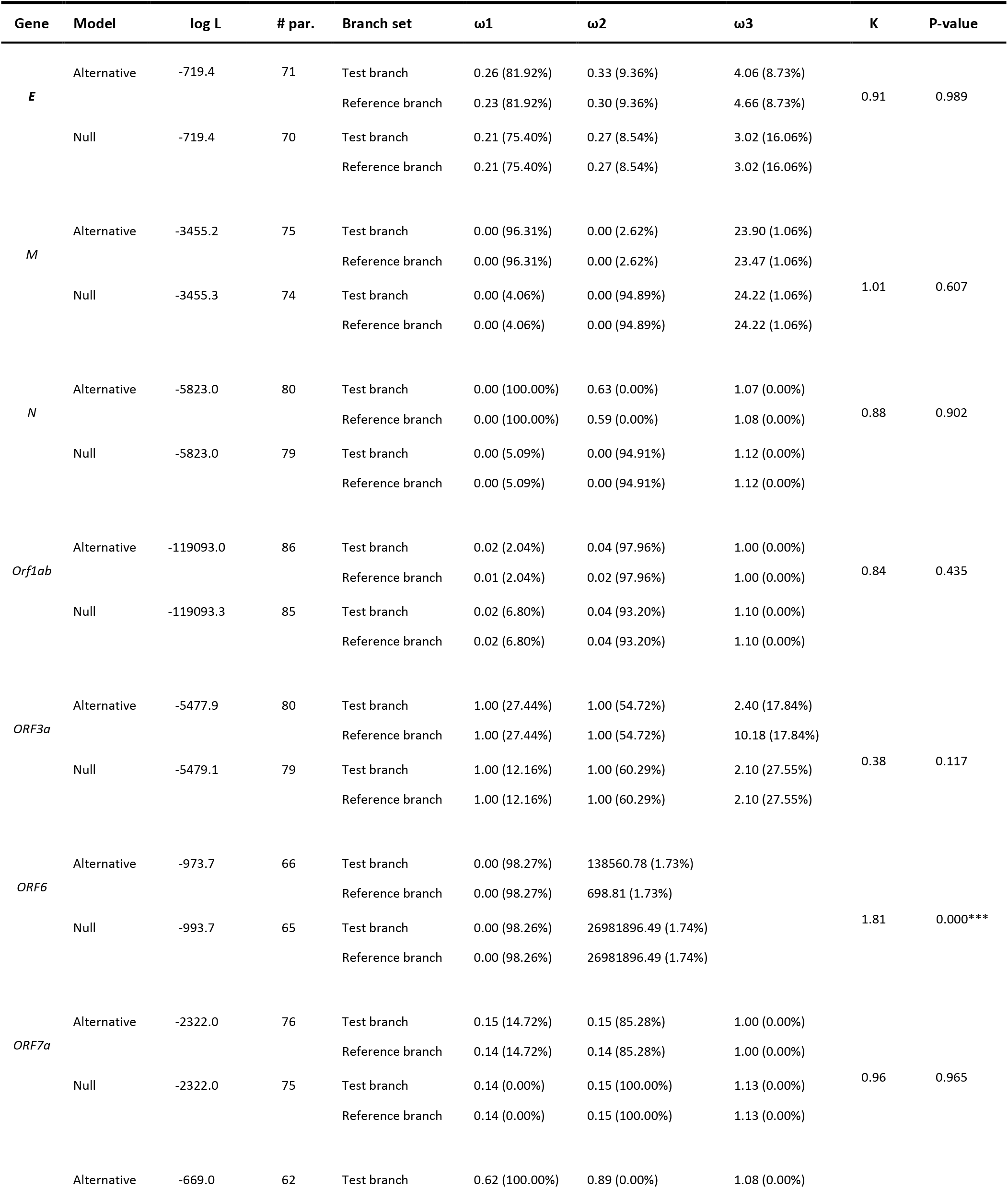
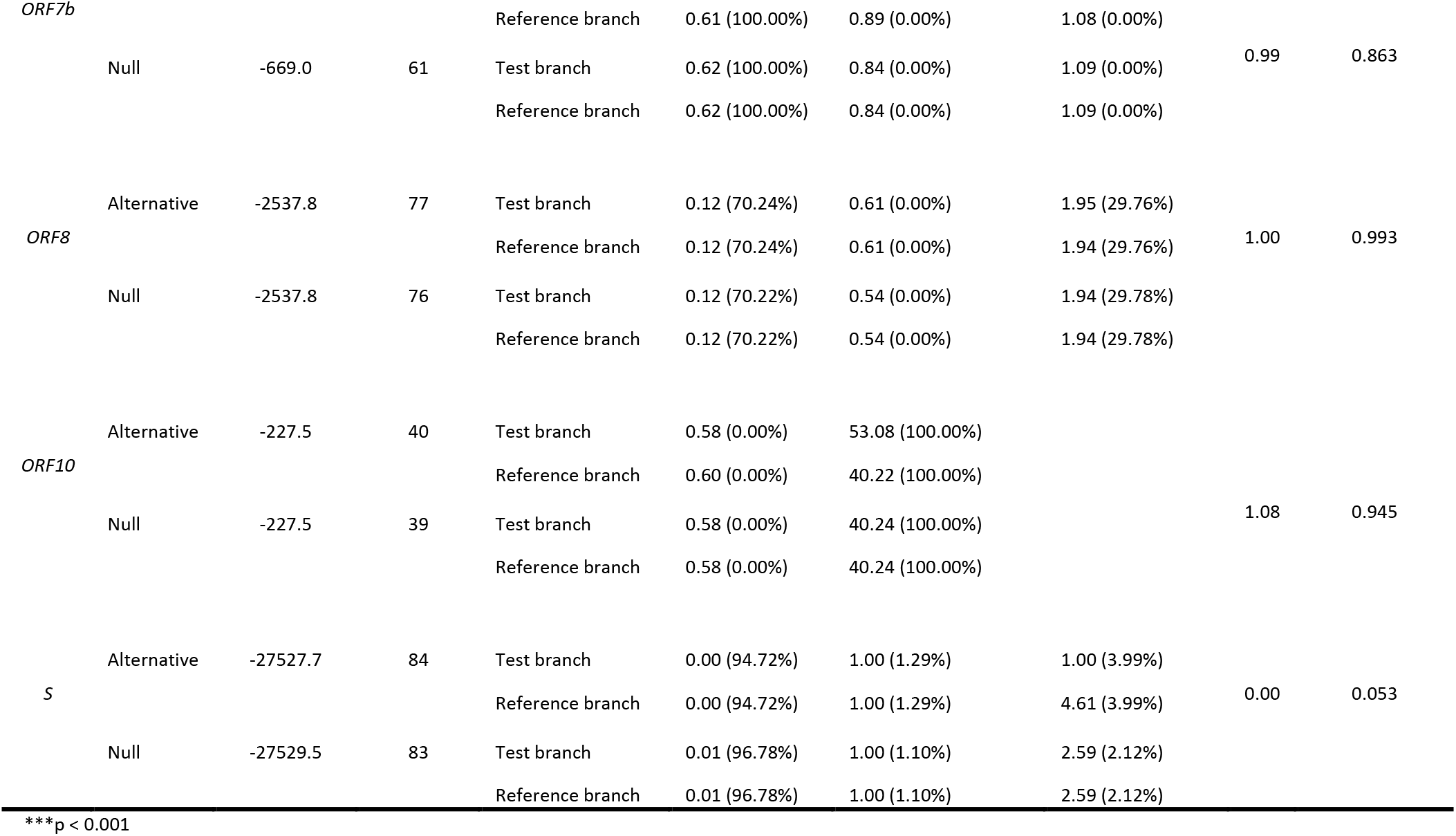
Selection intensity change of 11 genes along the SARS-CoV branch (test branch) relative to the common ancestral branch (reference branch) of SARS-CoV and its sister taxa (WIV16 and Rs4231).

**Table S3.** Branches under the positive selection of the *S* gene. Positive selections are analyzed using branch-site model. For convenience, only the ω values of the foreground branches are shown. Positive selection sites located in subunit 1 of the *S* gene are shown in grey. Underlining shows positive selected sites located in the RBD.

| Branch | Parameter estimates | 2ΔL | df | p-value | Positive selection sites |
| --- | --- | --- | --- | --- | --- |
| A | $p_0=0.901$ $p_1=0.069$ $p_{2a}=0.027$ $p_{2b}=0.002$<br>$\omega_0=0.030$ $\omega_1=1.000$ $\omega_{2a}=23.705$ $\omega_{2b}=23.705$ | 7.13 | 1 | 0.007 | 9F,292R,335Y,376H,393G,399Y,514M<br>529F,591Q,607S,673T,675S,774Y,820S,845V |
| B | $p_0=0.902$ $p_1=0.071$ $p_{2a}=0.024$ $p_{2b}=0.001$<br>$\omega_0=0.029$ $\omega_1=1.000$ $\omega_{2a}=49.941$ $\omega_{2b}=49.941$ | 32.59 | 1 | 1.139E-08 | 9F,11D,27L,63S,72A,110M,133F,174I<br>190V,192M,227S,228Q,234L,238T,240S<br>260E,458R,701S,733N, 810Q,999S,1135M |
| D | $p_0=0.917$ $p_1=0.066$ $p_{2a}=0.014$ $p_{2b}=0.001$<br>$\omega_0=0.030$ $\omega_1=1.000$ $\omega_{2a}=657.736$ $\omega_{2b}=657.736$ | 12.21 | 1 | 0.000 | 14R,25S,27L,39I,52R,68T,69T,85V<br>87N,125H,138H,176K, 216E |
| E | $p_0=0.922$ $p_1=0.067$ $p_{2a}=0.009$ $p_{2b}=0.000$<br>$\omega_0=0.030$ $\omega_1=1.000$ $\omega_{2a}=123.522$ $\omega_{2b}=123.522$ | 8.72 | 1 | 0.003 | 25S,27L,37P,52R,93F,149*<br>176K,,216E,438T,473F |
| F | $p_0=0.919$ $p_1=0.067$ $p_{2a}=0.011$ $p_{2b}=0.000$<br>$\omega_0=0.030$ $\omega_1=1.000$ $\omega_{2a}=108.595$ $\omega_{2b}=108.595$ | 29.62 | 1 | 5.267E-08 | 9F,27L,40P,56I,65L,110M,125H<br>139N,143V,161S,194L,224V,227S,376H |
| G | $p_0=0.925$ $p_1=0.069$ $p_{2a}=0.004$ $p_{2b}=0.000$<br>$\omega_0=0.030$ $\omega_1=1.000$ $\omega_{2a}=226.266$ $\omega_{2b}=226.266$ | 8.03 | 1 | 0.004 | 136V,144D,174I,181L,222D, 229D,376H<br>733N,756D |
| H | $p_0=0.915$ $p_1=0.069$ $p_{2a}=0.013$ $p_{2b}=0.001$<br>$\omega_0=0.030$ $\omega_1=1.000$ $\omega_{2a}=193.715$ $\omega_{2b}=193.715$ | 25.31 | 1 | 4.882E-07 | 55V,67N,69T,75F,128R,144D,161S<br>171L,177L,216E,276V,358K,408L |
| I | $p_0=0.917$ $p_1=0.069$ $p_{2a}=0.011$ $p_{2b}=0.000$<br>$\omega_0=0.030$ $\omega_1=1.000$ $\omega_{2a}=998.996$ $\omega_{2b}=998.996$ | 13.07 | 1 | 0.000 | 2S,48A,63S,73V,160F,209*,240S<br>259T,270L |
| J | $p_0=0.922$ $p_1=0.070$ $p_{2a}=0.005$ $p_{2b}=0.000$<br>$\omega_0=0.030$ $\omega_1=1.000$ $\omega_{2a}=999.000$ $\omega_{2b}=999.000$ | 9.72 | 1 | 0.001 | 75F,84I,112A, 128R,190V,191I |
| L | $p_0=0.926$ $p_1=0.071$ $p_{2a}=0.001$ $p_{2b}=0.000$<br>$\omega_0=0.030$ $\omega_1=1.000$ $\omega_{2a}=31.578$ $\omega_{2b}=31.578$ | 4.35 | 1 | 0.037 | 364T,380K,382Y,732I,955Y |

2 $\Delta$ L: twice difference of likelihood values between two nested models; df: degrees of freedom; proportion of sites and their corresponding $\omega$ values in four site classes ( $p_0$ , $p_1$ , $p_{2a}$ and $p_{2b}$ ) are shown. \* represents alignment gap in reference sequence.

